# Gut microbiota features associated with *Clostridioides difficile* colonization in dairy calves

**DOI:** 10.1101/2021.05.11.443551

**Authors:** Laurel E. Redding, Alexander S. Berry, Nagaraju Indugu, Elizabeth Huang, Daniel P. Beiting, Dipti Pitta

## Abstract

Diarrheal disease, a major cause of morbidity and mortality in dairy calves, is strongly associated with the health and composition of the gut microbiome. *Clostridioides difficile* is an opportunistic pathogen that proliferates and can produce enterotoxins when the host experiences gut dysbiosis. However, even asymptomatic colonization with *C. difficile* can be associated with differing degrees of microbiome disruption in a range of species, including people, swine, and dogs. Little is known about the interaction between *C. difficile* and the gut microbiome in dairy calves. In this study, we sought to define microbial features associated with *C. difficile* colonization in pre-weaned dairy calves less than 2 weeks of age. We characterized the fecal microbiota of 80 calves from 23 different farms using 16S rRNA sequencing and compared the microbiota of *C. difficile*-positive (n=24) and *C. difficile*-negative calves (n=56). Farm appeared to be the greatest source of variability in the gut microbiota. When controlling for calf age, diet, and farm location, there was no significant difference in Shannon alpha diversity (*P*= 0.50) or in weighted UniFrac beta diversity (P=0.19) between *C. difficile*-positive and –negative calves. However, there was a significant difference in beta diversity as assessed using Bray-Curtiss diversity (*P*=0.0077), and *C. difficile*-positive calves had significantly increased levels of *Ruminococcus (gnavus group)* (*Adj. P*=0.052)*, Lachnoclostridium* (*Adj. P*=0.060)*, Butyricicoccus* (*Adj. P*=0.060), and *Clostridium sensu stricto 2* compared to *C. difficile*-negative calves. Additionally, *C. difficile*-positive calves had fewer microbial co-occurrences than *C. difficile*–negative calves, indicating reduced bacterial synergies. Thus, while *C. difficile* colonization alone is not associated with dysbiosis and is therefore unlikely to result in an increased likelihood of diarrhea in dairy calves, it may be associated with a more disrupted microbiota.

## Introduction

Infectious diarrheal disease is one of the main causes of mortality in dairy calves (1, 2), and calves less than 30 days of age are at highest risk of developing diarrhea (3, 4). Studies have shown that gut microbial composition is associated with gut health and the likelihood of diarrhea: reductions in microbial diversity are associated with an increased incidence of diarrhea (5), and the colonization of the calf gut with beneficial bacteria along with the decreased colonization of potential pathogens decreases the likelihood of calf diarrhea (6).

*Clostridioides difficile* is a spore-forming anaerobic, gram-positive bacillus that is a significant enteric pathogen in many species of animals. Colonization with *C. difficile* has been shown to be associated with reduced gut microbial diversity and increased colonization of pathogenic bacteria in people (7, 8), and we recently demonstrated a similar association in puppies (9). Dairy calves, like the neonates of other species, are colonized with *C. difficile* at high rates, with reported prevalences ranging from 28-56% (10, 11). While there is some evidence that infection with *C. difficile* can result in diarrhea in calves (12), the effect of the asymptomatic colonization of calves on the gut microbiome is unknown. Given the crucial role of the gut microbiome in providing colonization resistance against pathogens that cause diarrhea (13, 14), a better understanding of the effect of pathogens such as *C. difficile* on the calf gut microbiome is needed. The goal of this study was thus to define the gut microbiota features associated with *C. difficile* colonization in dairy calves and to define the effects of calf age, diet, and farm on the risk of colonization.

## Methods

### Sample collection

Fecal samples were manually collected from up to five randomly selected healthy calves less than two weeks of age from each of 23 dairy farms in Pennsylvania, Maryland and Delaware. This study was approved by the Institutional Animal Care and Use Committee of the University of Pennsylvania.

### Detection of *C. difficile*

Individual fecal samples were tested for *C. difficile* using the Xpert *C. difficile* assay (Xpert CD assay; Cepheid, Sunnyvale, CA, USA) according to the manufacturer’s instructions. This assay detects the cytotoxin gene (*tcdB*) and binary toxin genes (*cdtA* and *cdtB*). Additionally, the assay has a callout for ribotype NAP1/B1/027.

To rule out the possibility of colonization with non-toxigenic *C. difficile*, pooled fecal samples from each farm were also submitted for anaerobic culture. Briefly, 0.5 g of formed fecal sample was mixed with 0.5 ml of 100% ethanol. The mixture remained for 60 minutes at room temperature before being inoculated on Cycloserine-cefoxitin fructose modified agar (CCFA) (Remel™) or *Clostridium difficile* Selective Agar (BBL™) and Columbia CNA agar (Thermo Fisher Scientific Remel Products). Inoculated plates and broth were incubated in BD Gas-Pak™ EZ container systems with BD BBL™ CO2 generators and BD BBL™ Gas Pak™ anaerobic CO2 indicators (Franklin Lakes, NJ) at 36°C ± 2°C under anaerobic growth conditions for seven days and checked for growth every other day. Suspect colonies were identified and isolated. Isolates were confirmed to be *C. difficile* by Maldi-TOF identification and/or RapID ANA II System (Thermo Fisher Scientific Remel Products).

### 16S rRNA sequencing

DNA was extracted from fecal samples using Qiagen PowerSoil DNA extraction kit. 16S rRNA sequencing was performed as described previously (9, 15). Briefly, the V4 region of the 16S rRNA gene was amplified using PCR, which was performed using Accuprime Pfx Supermix and custom primers for 30 cycles (15). PicoGreen quantification was used to normalize post-PCR products and AMPureXP beads were used to clean the combined pools. Libraries were quantified and sized using a Qubit 2.0 and Tapestation 4200, respectively. 250bp paired-end sequencing was performed using an Illumina MiSeq.

### Sequence data processing using QIIME2

The QIIME2 pipeline (16) was used to process and analyze 16S sequencing data. Samples were demultiplexed using q2-demux and denoised using Dada2 (17). Sequences were aligned using maaft (18) and phylogenetic trees were reconstructed using fasttree (19). Shannon alpha diversity, weighted UniFrac and Bray-Curtis beta diversity metrics were estimated using q2-core-metrics-diversity after samples were rarefied to 1941 reads per sample, and p-values were adjusted for multiple hypothesis testing using Benjamini-Hochberg (B-H) false discovery rate (FDR) corrections (20). Taxonomy was assigned to sequences using q2-feature-classifier classify-sklearn (21) against the Silva reference database (22). Taxa were collapsed to the genus level, when possible. OTUs with less than 1% average relative abundance across all samples were removed.

### Correlation analysis and differential feature selection

The correlation between *C. difficile* culture status and Shannon alpha diversity was determined using a linear mixed effects model as implemented in the lme4 package (23) in R where age was controlled for as a fixed effect and with farm and diet as random effects. The correlation between *C. difficile* culture status on gut microbiota beta diversity was determined using PERMANOVA as implemented in the vegan package (24) in R controlling for age, farm, and diet. Principal coordinate analyses were performed using the phyloseq package in R (25). Differentially-abundant taxa were determined using LDA Effect Size (LEfSe) (26) and Analysis of Composition of microbiomes (ANCOM), and p-values were adjusted for multiple hypothesis testing using B-H FDR corrections in R. The Dice index (27) was used to determine the co-occurrence of bacterial genera. Boxplots and LEfSe plots were visualized using ggplot2 (28) and ggthemes.

## Results

### Subject characteristics and *C. difficile* status

Fecal samples were collected from a total of 92 Holstein calves from 23 farms. All calves appeared systemically healthy at the time of sampling and none had received antimicrobial therapy. The mean (SD) age of the calves was 7.0 (5.0) days. Thirty-six (35.6%) calves were fed waste milk, while the remaining calves were fed either colostrum or whole milk.

*C. difficile* was detected by qPCR in 28 calves (30.4%, 95% CI 21.2-40.9%) (**Fig. 1**). Of the 28 samples that were positive for *C. difficile* on qPCR, 1 (3.6%) was positive for Toxin B only, 14 (50%) were positive for binary toxin only, and 13 (46.4%) were positive for both Toxin B and the binary toxin. None of the organisms were identified as the NAP1/B1/027 ribotype. On 14 farms, there were both *C. difficile*-positive and *C. difficile*-negative calves, whereas on the remaining farms, all of the calves were *C. difficile*-negative. There were no farms where all samples were qPCR-negative but the pooled sample was culture-positive. Neither calf age nor feeding of waste milk were significantly associated with the likelihood of detecting *C. difficile* among the calves (OR=1.01, p=0.805 and OR=0.71, p=0.493, respectively) (**Fig. 1**).

**Figure 1:**
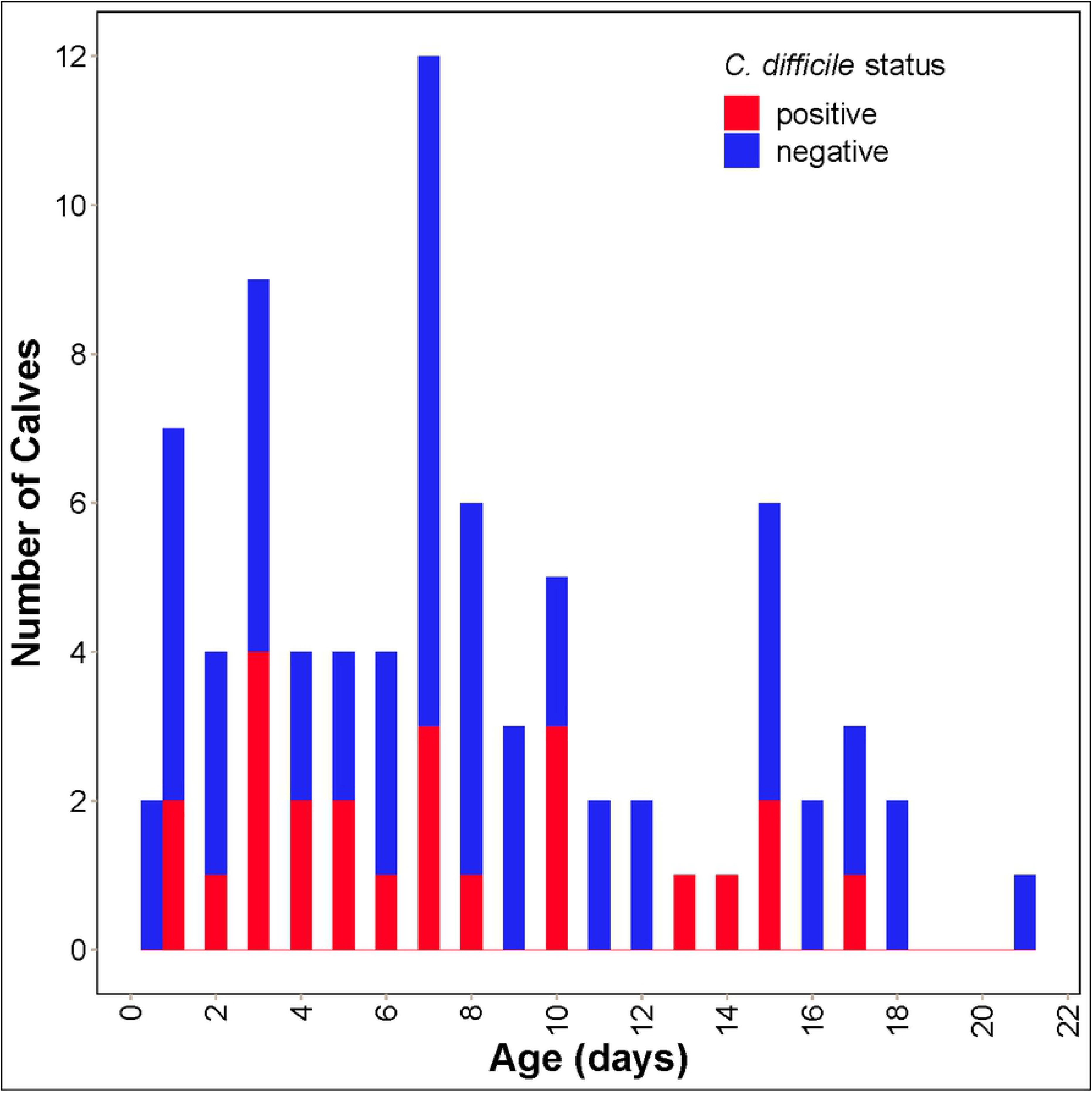
Distribution of age and *C. difficile* colonization status in 92 pre-weaned Holstein dairy calves

### Effect of *C. difficile* status on microbiota diversity

Microbiota community structure of 87 calf fecal samples was assessed by sequencing and analyzing the V4 region of the 16S rRNA gene. Three samples were dropped from subsequent analyses because of low coverage and four additional samples were dropped because there was not enough sample for qPCR analysis. Among the 80 remaining samples, 24 were positive for *C. difficile* by qPCR and 56 were negative (**Fig 1**).

The relationship between *C. difficile* infection and microbial diversity of the gut microbiota was assessed. Since calves ranged in age, diet, and farm location, a linear mixed effects model was performed to assess the relationship between *C. difficile* infection and alpha diversity by setting age as a fixed variable and farm and feeding type as random-effect variables. The association between *C. difficile* status and Shannon alpha diversity was not significant (*P*= 0.50) as determined by ANOVA when controlling for age, diet, and farm location (**Fig. 2**). PERMANOVA was then used to test associations between *C. difficile* infection status and beta diversity of the gut microbiome. Farm location alone explained most of the variation in gut microbiota composition across samples using both Bray-Curtis (*P*=1e-4; R2=0.43) and weighted UniFrac (*P*=1e-4; R2=0.46) beta diversity metrics (**Fig. 3, Fig. 4**). Age and diet were not significantly associated with gut microbiota composition after controlling for farm (*P*>0.1). After controlling for farm, age, and diet, *C. difficile* status was significantly associated with Bray-Curtis beta diversity (*P*=0.0077; R2=0.023), explaining 2.3% of the variation in gut microbiota composition. *C. difficile* status was not significantly associated with weighted UniFrac beta diversity (*P*= 0.1934; R2=0.013) after controlling for farm, age, and diet (**Fig. 3**). Some clustering by farm and by *C. difficile* status within farms was apparent on principal coordinate analysis (**Fig. 4**).

**Figure 2:**
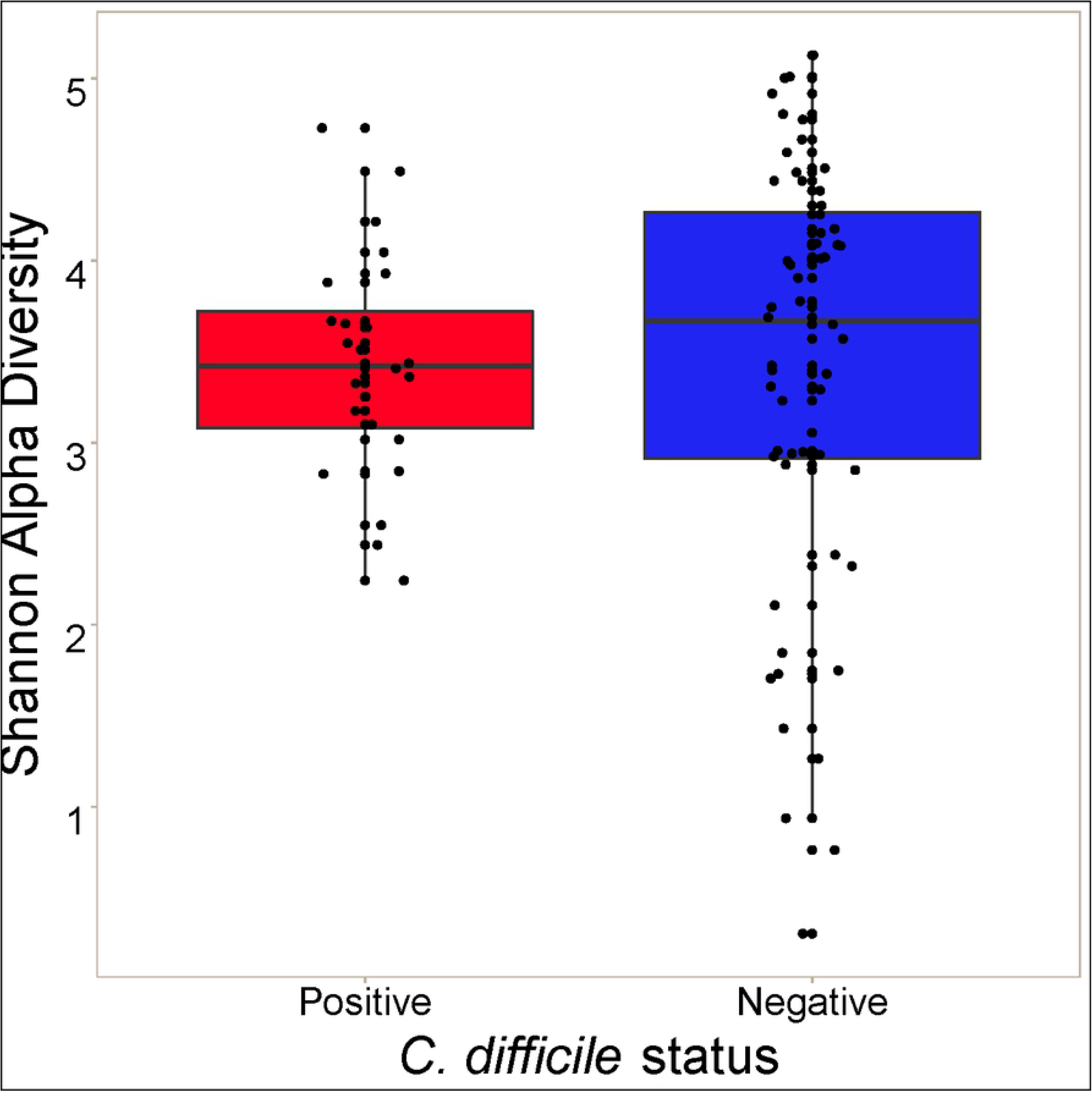
Alpha diversity of the gut microbiome in 86 pre-weaned Holstein dairy calves by *C. difficile* colonization status

**Figure 3:**
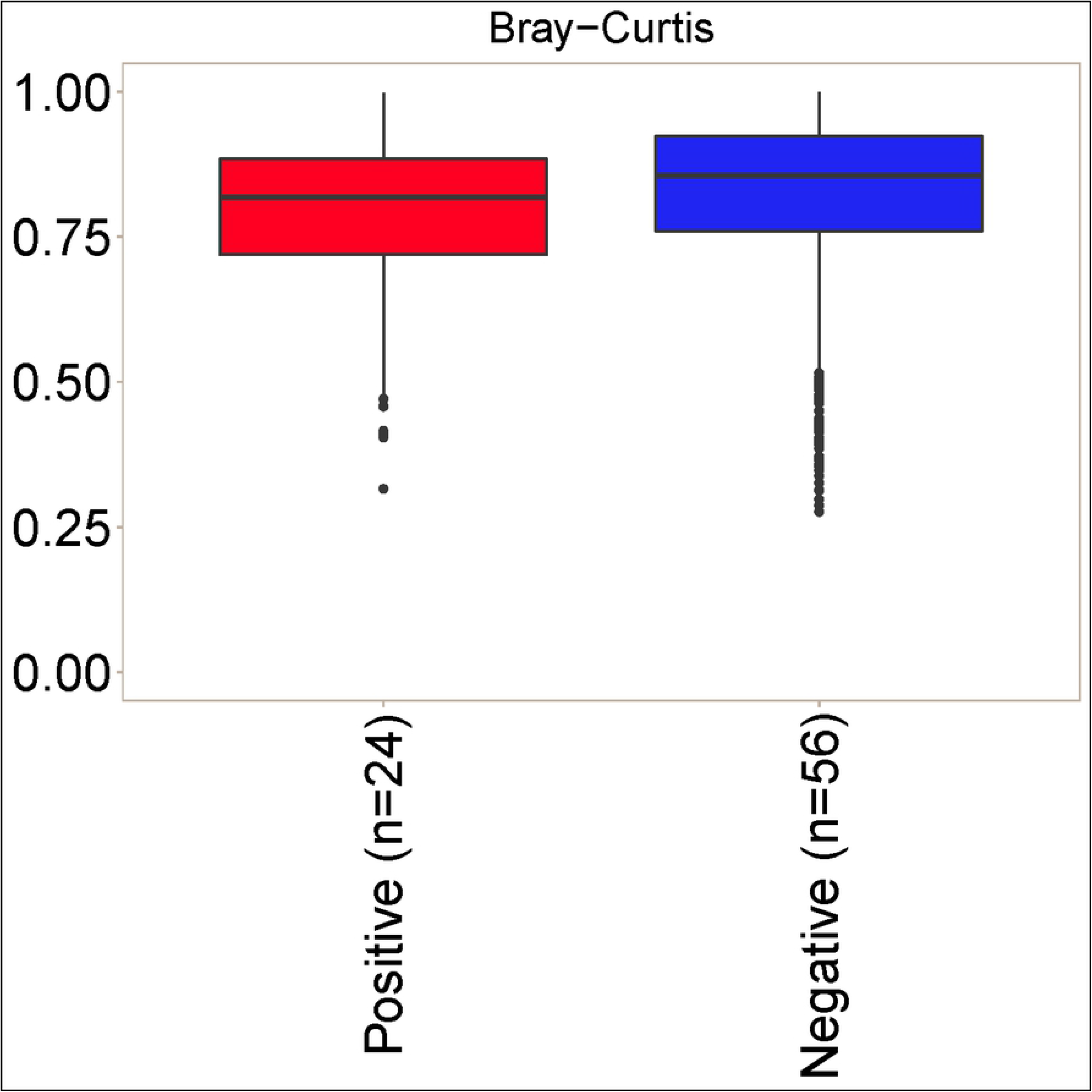
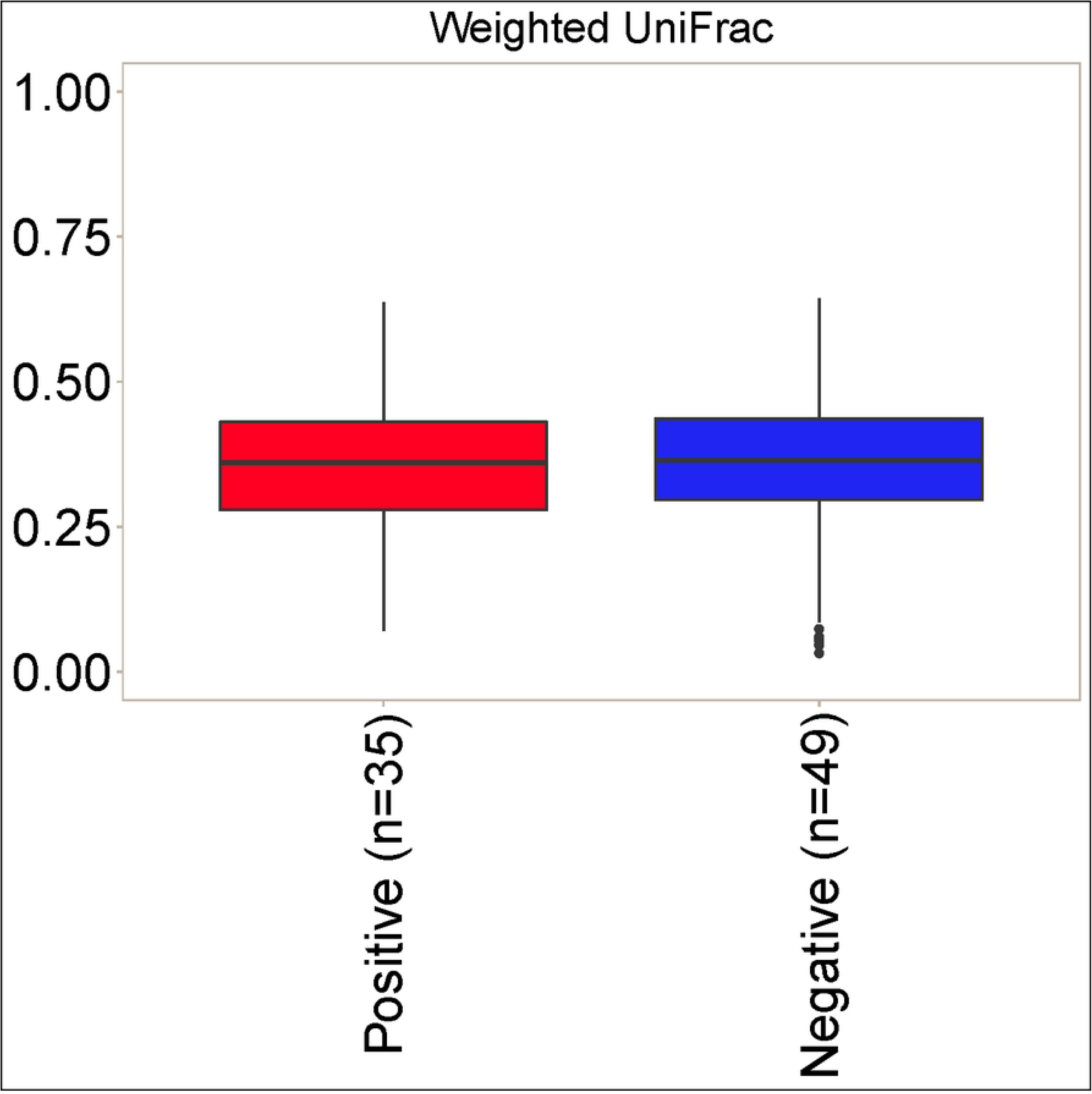
Beta diversity of the gut microbiome in 86 pre-weaned Holstein dairy calves by *C. difficile* colonization status. A. Bray-Curtis beta diversity. B. Weighted UniFrac.

**Figure 4:**
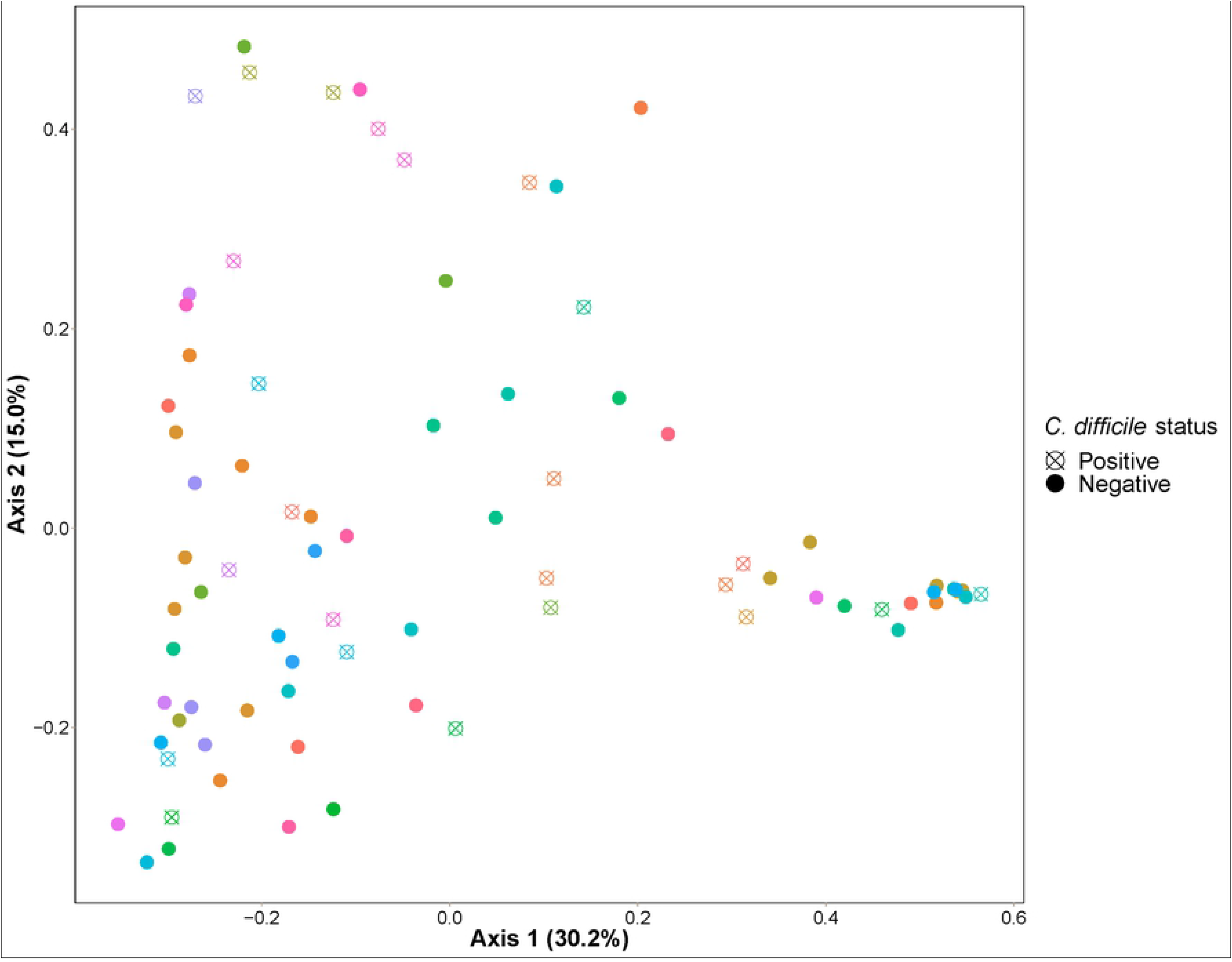
Bray-Curtis principal coordinate analysis (PCoA) of fecal samples from 86 pre-weaned dairy calves by *C. difficile* colonization status and by farm

### Bacterial community composition

Since *C. difficile* status was associated with differences in gut microbiota composition as determined by beta diversity, we next sought to determine the specific bacterial taxa associated with *C. difficile* infection. At the phylum level, there were no significant differences between bacterial communities in *C. difficile*-positive and -negative samples (**Fig. 5**). The Firmicutes phylum predominated (57.1% in *C. difficile*-positive samples and 51.4% in *C. difficile*-negative samples), followed by Proteobacteria (17.1% and 24.3%), Bacteroides (16.7% and 11.5%), and Actinobacteria (8.1% and 9.7%).

**Figure 5:**
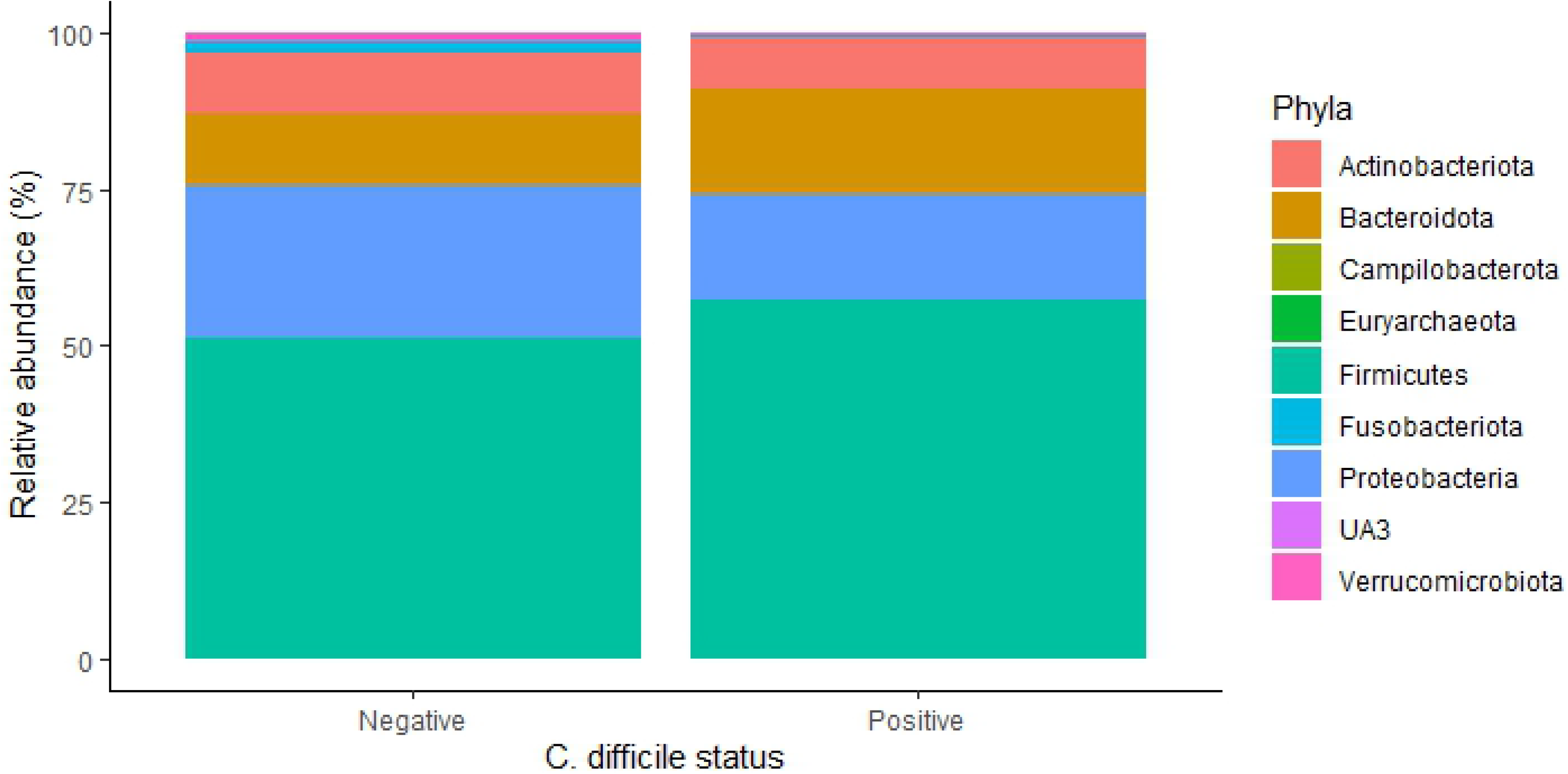
Distribution of bacterial phyla by *C. difficile* status in fecal samples from 86 pre-weaned dairy calves. The nine most abundant phyla are displayed.

At the genus level, the only significant difference between *C. difficile*-positive and –negative samples by ANCOM occurred for Clostridioides. When considering LEFse analysis, there were four taxa among the 19 taxa with average relative abundance greater than 1% that were statistically significantly (*Adj. P*<0.1) associated with *C. difficile* status. *Ruminococcus (gnavus group)* (*Adj. P*=0.052)*, Lachnoclostridium* (*Adj. P*=0.060)*, Butyricicoccus* (*Adj. P*=0.060), and *Clostridium* (*sensu stricto 2*) (*Adj. P*=0.064) were all found in higher abundance among *C. difficile-*positive calves than in *C. difficile-*negative calves (**Fig. 6**). While not statistically significantly different among the two groups, levels of *Lactobacillus, Megasphaera*, and *Streptococcus* were increased in *C. difficile*-positive samples, while levels of *Blautia, Fusobacterium, Tyzzerella, Enterobacteriaceae, Fecalibacterium, Dorea*, and *Collinsella* were decreased.

**Figure 6:**
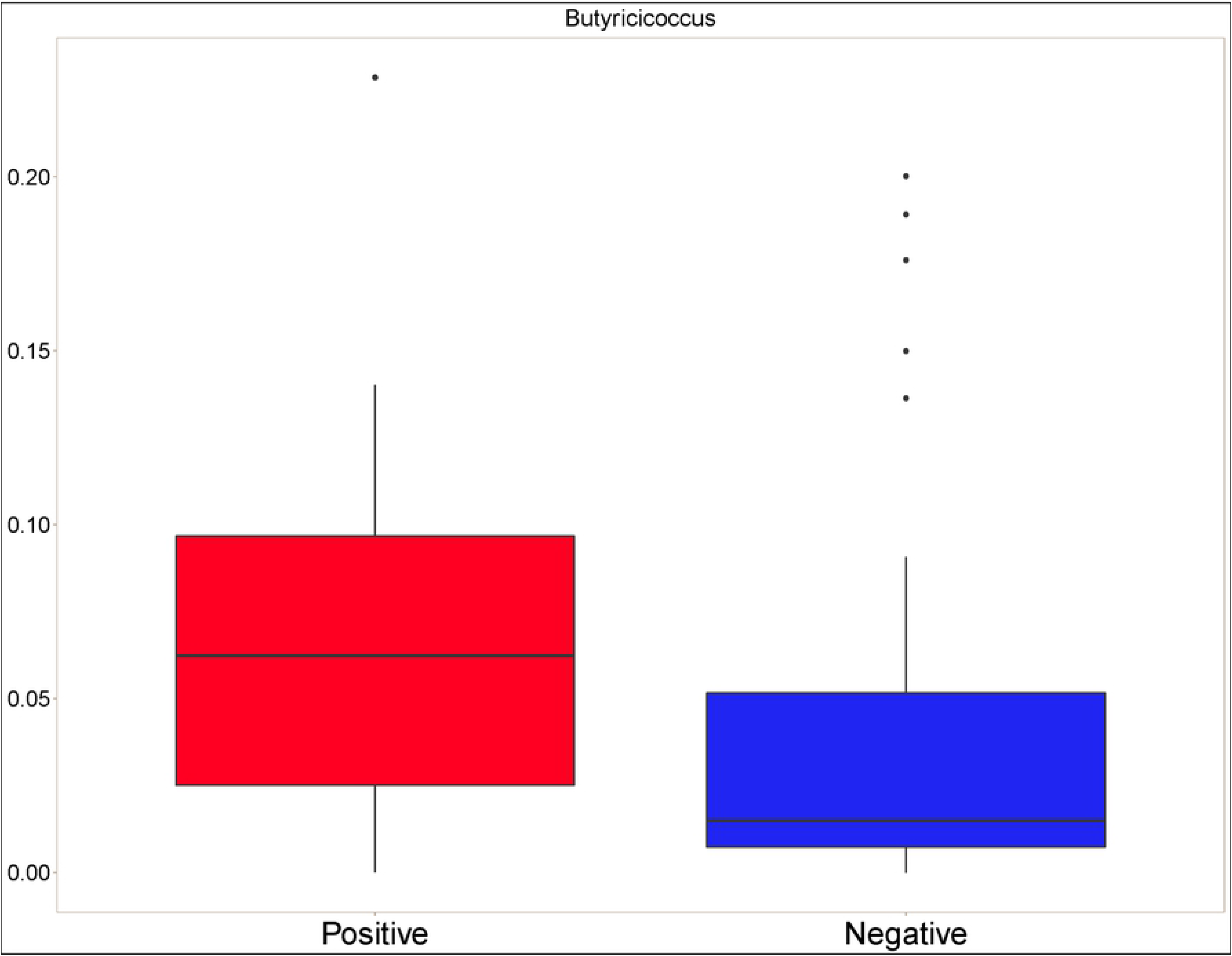
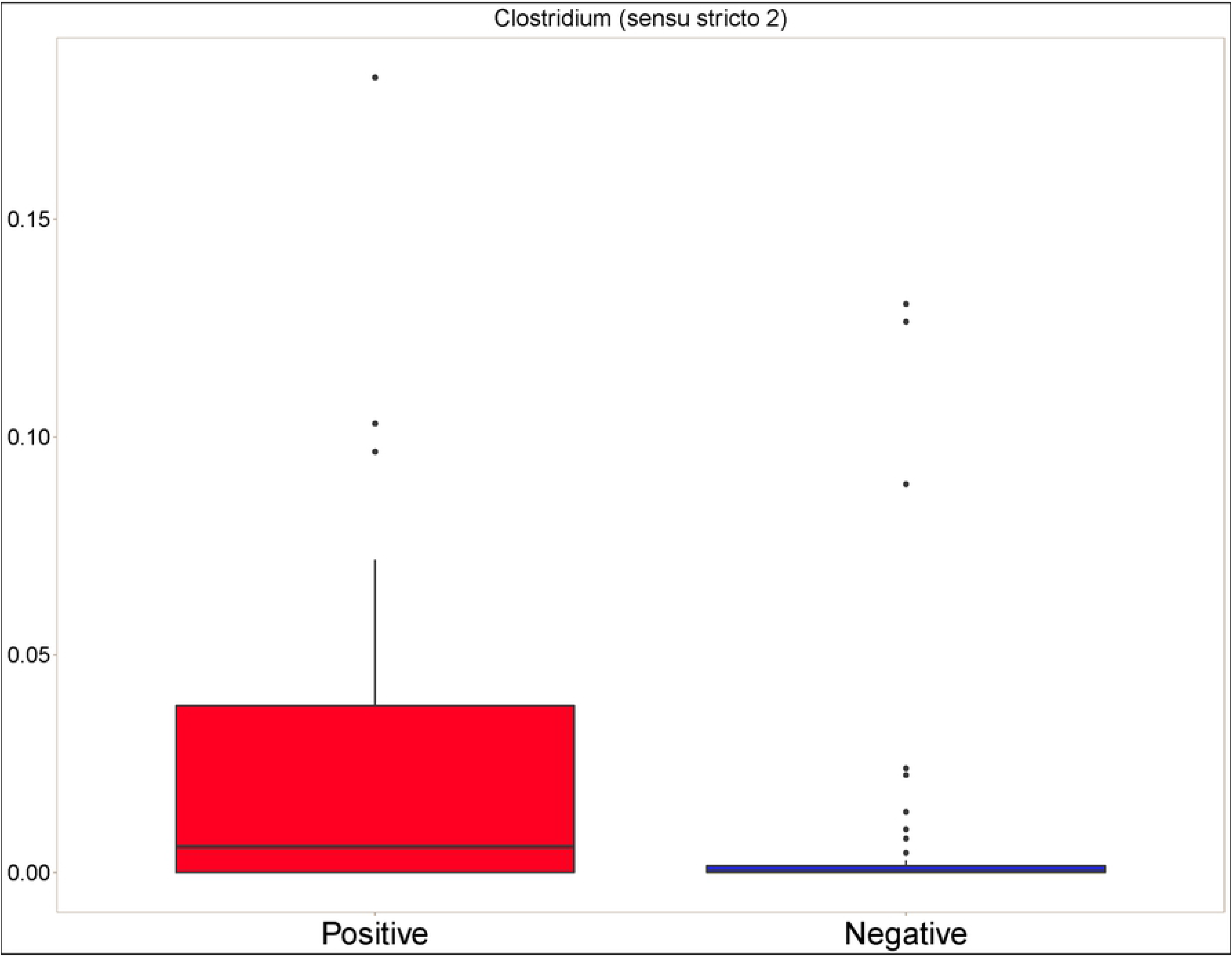
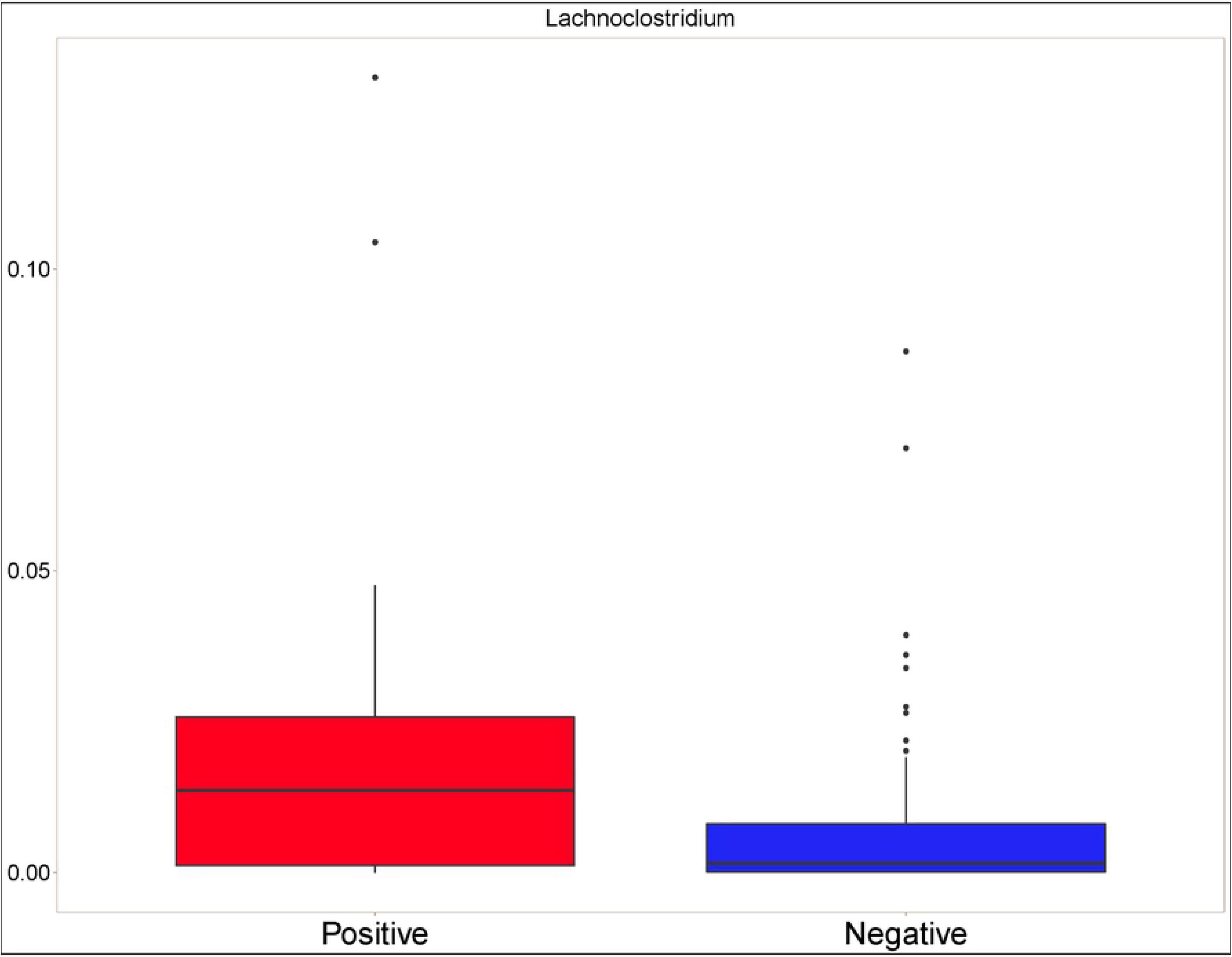
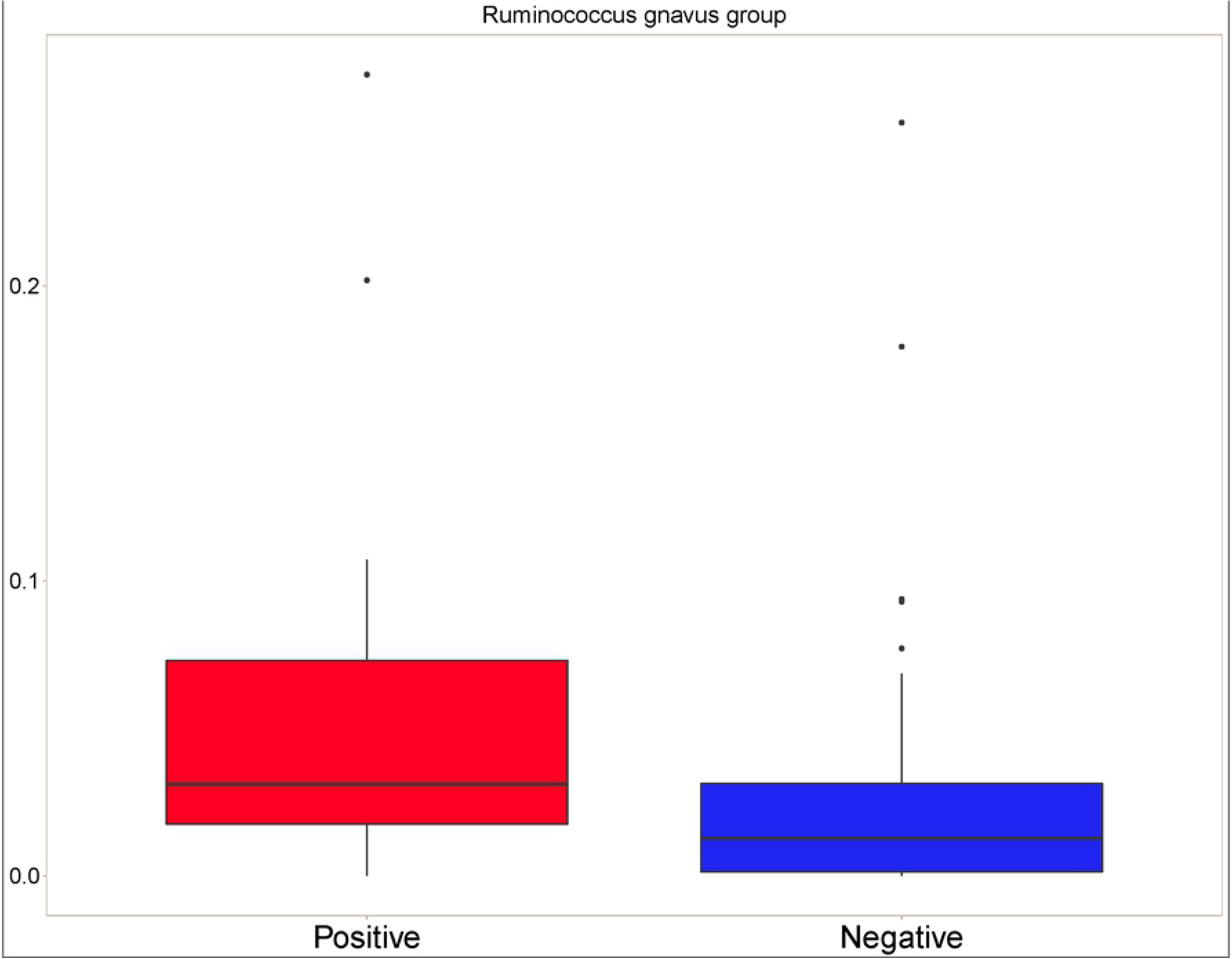
Distribution of bacterial taxa that were found at higher levels in *C. difficile*-positive calves by *C. difficile* colonization status in 86 pre-weaned Holstein dairy calves. A. *Butyricicoccus*. B. *Clostridium sensu stricto 2*. C. *Ruminococcus gnavus*. D. *Lachnoclostridium*.

Because microbes work synergistically in the gut, we sought to determine the associative interactions between bacteria using a co-occurrence analysis based on the Dice index. When considering all levels of abundance, more co-occurrence of bacterial taxa appeared in the *C. difficile*-negative samples, with 1,488 (65.5%) highly (correlation coefficient>0.6) and significantly (p<0.01) correlated genera pairs. Most co-occurrences were among members of the Firmicutes phylum (1295, 55.0%). However, members of Firmicutes also showed high co-occurrence with Actinobacteria and Bacteroidetes. In the *C. difficile*-positive samples, there were fewer highly co-occurring genera, with 830 (73.3%) highly and significantly correlated genera pairs. When only considering taxa with levels of abundance greater than 1%, there were no significant differences in co-occurrence patterns **(Fig. 7)**.

**Figure 7.**
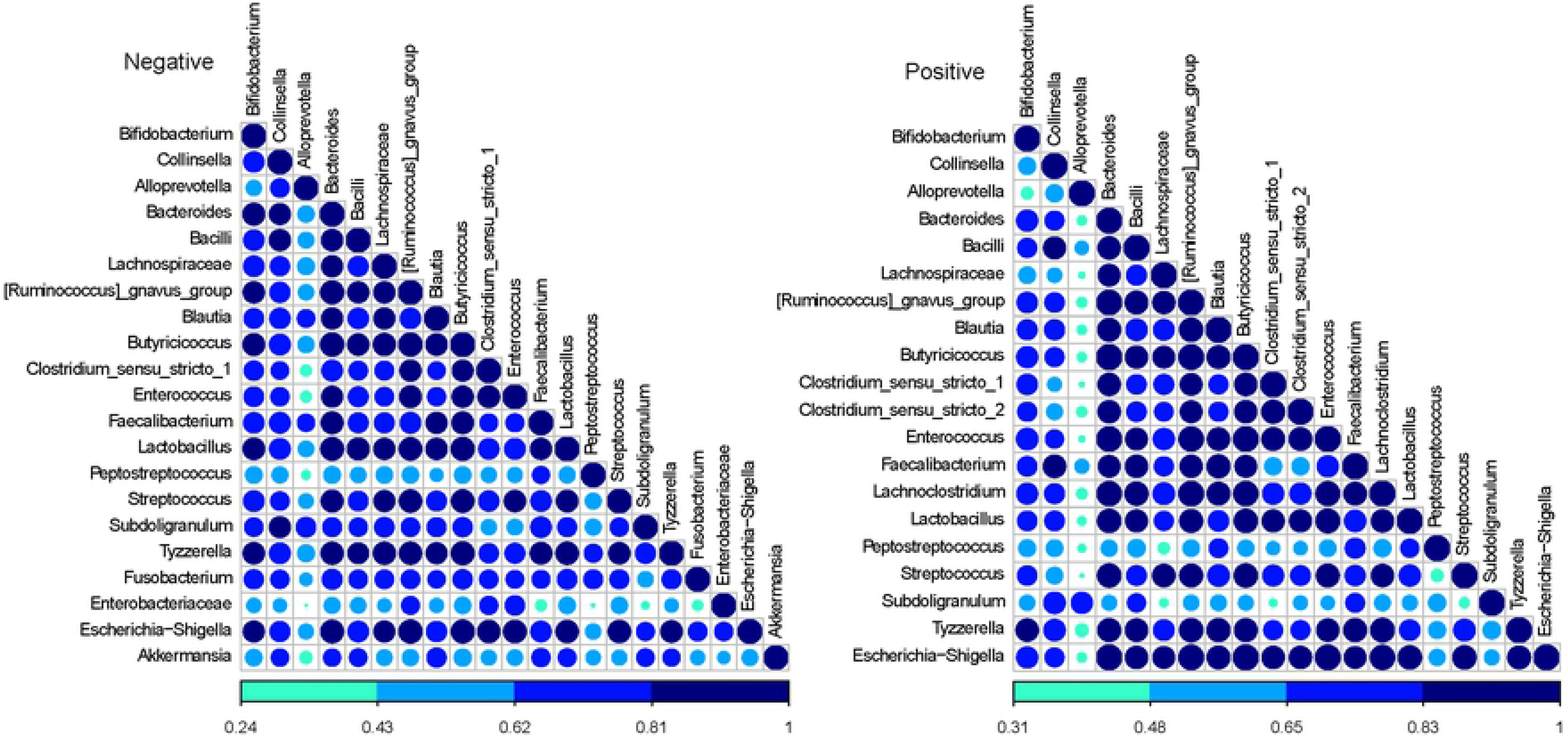
Analysis of co-occurrence among microbial lineages scored using the Dice index by *C. difficile*-colonization status (positive and negative). Dice indexes are shown as a heat map for all genera present at a level of abundance greater than 1% and with statistically significant (p<0.01) co-occurrence are shown as a heatmap. The degree of co-occurrence is shown by the color code at the bottom.

## Discussion

In this study, we characterized microbial features associated with asymptomatic *C. difficile* colonization in dairy calves. While the role of *C. difficile* in calf diarrhea remains equivocal (12), exploring the association between this pathogen and the gut microbiome is important for understanding factors that affect gut health and enteric diseases. While a number of studies have examined the epidemiology of *C. difficile* in animals of veterinary importance, the association between the microbiome and *C. difficile* is only beginning to be explored in dogs (9), horses (29), and pigs (30). Notably, in pigs, the presence of *C. difficile* is associated with significantly reduced microbial diversity and increased levels of enteropathogens associated with neonatal diarrhea (30).

Unsurprisingly, as in other studies (31–33), we found that the farm was the source of most of the variation in gut microbiota composition. However, even among calves from the same farm, there was variability in both *C. difficile* colonization status and gut microbial diversity, suggesting, as have other studies (32, 34), that the farm environment is only one of many competing influencers of the developing calf gut microbiome. Neither diet nor age were significantly associated with microbiome composition when controlling for farm, but this is almost certainly due to the small sample size within each farm and the lack of within-farm variability in factors such as diet. When controlling for age, diet, and farm, we noted a significant difference in beta diversity between *C. difficile*-positive and *C. difficile*-negative fecal samples when considering the Bray-Curtis metric but not the unweighted UniFrac metric. While both of these metrics are weighted by abundance, the latter metric weighs diversity by phylogenetic relationship. Thus the lack of a significant difference when considering the weighted UniFrac metric suggests that, while there may be a significant difference in the composition of microbial communities, the differentially-abundant microbes might be closely related to one another. Indeed, all four genera identified as differentially-abundant by LEfSe are members of the *Clostridia* class, with two belonging to the *Clostrideaceae* family.

While the lack of a consistent difference in alpha and beta diversity between *C. difficile*-positive and *C. difficile*-negative samples suggests that the effect of *C. difficile* colonization on the gut microbiome of calves is minimal, other findings suggest that *C. difficile* colonization is associated with a more disrupted – but not dysbiotic – gut microbiome. *C. difficile* colonization was preferentially associated with certain bacterial taxa of the class *Clostridia* that do have associations with dysbiosis. Notably, the overrepresentation of *Ruminococcus gnavus* and *Lachnoclostridia* in *C. difficile*-positive calves point to the possibility of an underlying imbalance in the gut microbiome. *R. gnavus,* a Gram-positive anaerobe that is typically found in the gut of over 90% of healthy people at abundances less than 0.1%, has been robustly associated with inflammatory dysbiotic conditions such as Crohn’s disease (35–37), allergic airway disease (38), eczema (39), and spondyloarthritis (40). Dramatic blooms of *R. gnavus* occur in patients experiencing flares of inflammatory bowel disease, with abundance levels that can peak at 69% of the gut microbiota (37). Notably, this association appears to occur across species, as the gut microbiomes of both infants (7) and piglets (30) colonized with *C. difficile* also had increased levels of *Ruminococcus* species, including *R. gnavus*. Additionally, *Ruminococcus* was one of six bacterial genera in the gut microbiome that predicted the occurrence of diarrhea in calves in another study (41). The increased relative abundance of *Clostridium sensu stricto* and *Lachnoclostridia* in *C. difficile*-positive calves also points to the possibility of a less healthy gut environment. An increased relative abundance of *Clostridium sensu stricto*, which was also found in *C. difficile*-positive piglets (30), was associated with food allergies in infants (42) and diarrhea in piglets (43). A tentative association between increased levels of *Lachnoclostridia* and neoplasia of the gastrointestinal tract has been identified in people (44, 45). While no such association has been explored in animals, the overrepresentation of this taxon in *C. difficile*-positive calves may be the result of a more disrupted gut microbiota. However, it is also important to note that the increased relative abundance of these taxa were only detected using LEfSe analysis and not ANCOM, which suggests that the association is likely relatively weak.

Certain bacterial taxa that predominate in healthy calves were found at lower (but not statistically significantly lower) levels in *C. difficile*-positive calves. Notably, *Fecalibacterium, Dorea, Enterobacteriaceae* and *Collinsella* are among the most abundant genera in healthy pre-weaned calves (46–49), and some of these taxa provide colonization resistance against *C. difficile* (8, 50). Their decreased relative abundance in *C. difficile*-positive calves is thus also reflective of a more disrupted gut microbiome. The decreased co-occurrence of bacterial taxa in *C. difficile*-positive calves compared to *C. difficile*-negative calves when considering all levels of abundance may also corroborate the notion of a slightly more disrupted gut microbiome in colonized calves. However, because the difference occurred only in rare taxa (abundance < 1%), this difference appears unlikely to result in dysbiosis.

One finding that is in contradiction to the general trend of *C. difficile* colonization being associated with disrupted microbiota is the increased abundance of *Butyricicoccus* in *C. difficile*-positive calves. In people, *Butyricicoccus* species of bacteria are generally found in *lower* levels in people colonized with *C. difficile* (51) or diagnosed with inflammatory bowel disease (52, 53), and at higher levels in healthy dairy calves compared to calves with diarrhea (48, 54). It is unclear why they were found at higher levels in *C. difficile*-positive calves compared to *C. difficile*-negative calves. *Butyricicoccus* bacteria produce butyrate, an important nutrient source for gut colonocytes and a beneficial driver of the immunological maturation of the gut mucosa (55), and account for one of the most abundant genera in dairy calves 7 days after birth (56). The differential levels in calves compared to people with enteric disease may be due to species-specific patterns of development of the neonatal gut. Species-specific differences may also explain why *C. difficile* colonized calves had higher levels of *Clostridial* genuses but colonized puppies had lower levels (9). While rumen development is minimal in pre-weaned calves, they are nevertheless ruminants and thus have fundamentally different enteric physiologies and microbial ecologies compared to true monogastric species.

Some limitations apply to this study. Heterogeneity in farm location, age, and diet across all of the sampled calves may have obscured features of the microbiome that would otherwise have been associated with *C. difficile* colonization. The cross-sectional nature of the study also precludes the possibility of drawing any conclusions about the duration of colonization and its effect on an already rapidly evolving gut microbiome. Finally, because we used qPCR to detect *C. difficile* in the calves’ feces, we were unable to detect non-toxigenic *C. difficile*. It is likely that toxigenic and non-toxigenic *C. difficile* occupy a similar ecological niche and compete for similar resources within the gut microbiota; thus the presence of non-toxigenic *C. difficile* could account for the lack of a significant difference in alpha diversity and microbial composition between *C. difficile*-positive and *C. difficile*-negative calves. However, we believe this possibility to be unlikely, as there were no samples that were negative on qPCR but came from a farm where the pooled sample was positive for *C. difficile* on anaerobic culture.

## Conclusion

The greatest source of variability in the calf microbiome was the farm, and there were few or no statistically significant differences in alpha or beta diversity between *C. difficile*-positive and *C. difficile*-negative calves. *C. difficile* colonization thus does not appear to be associated with dysbiosis or with increased levels of enteropathogens that cause calf diarrhea. However, microbial community signatures – including increased relative abundance of bacterial taxa that that have been associated with dysbiotic states in other species and in people - suggest that the microbiota of *C. difficile*-colonized calves is more disrupted than that of non-colonized calves.

## REFERENCES

1. McGuirk SM. Disease management of dairy calves and heifers. Vet Clin North Am Food Anim Pract. 2008 Mar;24(1):139–53. PubMed PMID: 18299036.

2. Foster DM, Smith GW. Pathophysiology of diarrhea in calves. Vet Clin North Am Food Anim Pract. 2009 Mar;25(1):13–36, xi. PubMed PMID: 19174281.

3. Virtala AM, Mechor GD, Grohn YT, Erb HN. Morbidity from nonrespiratory diseases and mortality in dairy heifers during the first three months of life. Journal of the American Veterinary Medical Association. 1996 Jun 15;208(12):2043–6. PubMed PMID: 8707681.

4. Gulliksen SM, Jor E, Lie KI, Hamnes IS, Loken T, Akerstedt J, et al. Enteropathogens and risk factors for diarrhea in Norwegian dairy calves. J Dairy Sci. 2009 Oct;92(10):5057–66. PubMed PMID: 19762824.

5. Oikonomou G, Teixeira AG, Foditsch C, Bicalho ML, Machado VS, Bicalho RC. Fecal microbial diversity in pre-weaned dairy calves as described by pyrosequencing of metagenomic 16S rDNA. Associations of Faecalibacterium species with health and growth. PloS one. 2013;8(4):e63157. PubMed PMID: 23646192. Pubmed Central PMCID: 3639981.

6. Malmuthuge N, Chen Y, Liang G, Goonewardene LA, Guan le L. Heat-treated colostrum feeding promotes beneficial bacteria colonization in the small intestine of neonatal calves. J Dairy Sci. 2015 Nov;98(11):8044–53. PubMed PMID: 26342981.

7. Rousseau C, Levenez F, Fouqueray C, Dore J, Collignon A, Lepage P. Clostridium difficile colonization in early infancy is accompanied by changes in intestinal microbiota composition. J Clin Microbiol. 2011 Mar;49(3):858–65. PubMed PMID: 21177896. Pubmed Central PMCID: PMC3067754.

8. Zhang L, Dong D, Jiang C, Li Z, Wang X, Peng Y. Insight into alteration of gut microbiota in Clostridium difficile infection and asymptomatic C. difficile colonization. Anaerobe. 2015 Aug;34:1–7. PubMed PMID: 25817005.

9. Berry ASF, Kelly BJ, Barnhart D, Kelly DJ, Beiting DP, Baldassano RN, et al. Gut microbiota features associated with Clostridioides difficile colonization in puppies. PloS one. 2019;14(8):e0215497. PubMed PMID: 31469837. Pubmed Central PMCID: PMC6716646.

10. Houser BA, Soehnlen MK, Wolfgang DR, Lysczek HR, Burns CM, Jayarao BM. Prevalence of Clostridium difficile toxin genes in the feces of veal calves and incidence of ground veal contamination. Foodborne Pathog Dis. 2012 Jan;9(1):32–6. PubMed PMID: 21988399.

11. Knight DR, Thean S, Putsathit P, Fenwick S, Riley TV. Cross-sectional study reveals high prevalence of Clostridium difficile non-PCR ribotype 078 strains in Australian veal calves at slaughter. Appl Environ Microbiol. 2013 Apr;79(8):2630–5. PubMed PMID: 23396338. Pubmed Central PMCID: 3623178.

12. Hammitt MC, Bueschel DM, Keel MK, Glock RD, Cuneo P, DeYoung DW, et al. A possible role for Clostridium difficile in the etiology of calf enteritis. Veterinary microbiology. 2008 Mar 18;127(3-4):343–52. PubMed PMID: 17964088.

13. Malmuthuge N, Griebel PJ, Guan le L. The Gut Microbiome and Its Potential Role in the Development and Function of Newborn Calf Gastrointestinal Tract. Front Vet Sci. 2015;2:36. PubMed PMID: 26664965. Pubmed Central PMCID: PMC4672224.

14. Malmuthuge N, Guan LL. Understanding the gut microbiome of dairy calves: Opportunities to improve early-life gut health. J Dairy Sci. 2017 Jul;100(7):5996–6005. PubMed PMID: 28501408.

15. Kozich JJ, Westcott SL, Baxter NT, Highlander SK, Schloss PD. Development of a dual-index sequencing strategy and curation pipeline for analyzing amplicon sequence data on the MiSeq Illumina sequencing platform. Appl Environ Microbiol. 2013 Sep;79(17):5112–20. PubMed PMID: 23793624. Pubmed Central PMCID: PMC3753973.

16. Bolyen E, Rideout JR, Dillon MR, Bokulich NA, Abnet CC, Al-Ghalith GA, et al. Reproducible, interactive, scalable and extensible microbiome data science using QIIME 2. Nature Biotechnology. 2019 2019/08/01;37(8):852–7.

17. Callahan BJ, McMurdie PJ, Rosen MJ, Han AW, Johnson AJ, Holmes SP. DADA2: High-resolution sample inference from Illumina amplicon data. Nat Methods. 2016 Jul;13(7):581–3. PubMed PMID: 27214047. Pubmed Central PMCID: 4927377.

18. Katoh K, Standley DM. MAFFT multiple sequence alignment software version 7: improvements in performance and usability. Mol Biol Evol. 2013 Apr;30(4):772–80. PubMed PMID: 23329690. Pubmed Central PMCID: PMC3603318.

19. Price MN, Dehal PS, Arkin AP. FastTree 2#x002D;#x002D;approximately maximum-likelihood trees for large alignments. PloS one. 2010 Mar 10;5(3):e9490. PubMed PMID: 20224823. Pubmed Central PMCID: 2835736.

20. Benjamini Y, Hochberg Y. Controlling the False Discovery Rate: A Practical and Powerful Approach to Multiple Testing. Journal of the Royal Statistical Society Series B (Methodological). 1995;57(1):289–300.

21. Bokulich NA, Kaehler BD, Rideout JR, Dillon M, Bolyen E, Knight R, et al. Optimizing taxonomic classification of marker-gene amplicon sequences with QIIME 2’s q2-feature-classifier plugin. Microbiome. 2018 2018/05/17;6(1):90.

22. Quast C, Pruesse E, Yilmaz P, Gerken J, Schweer T, Yarza P, et al. The SILVA ribosomal RNA gene database project: improved data processing and web-based tools. Nucleic Acids Res. 2013;41(Database issue):D590–D6. PubMed PMID: 23193283. Epub 11/28. eng.

23. Bates D, Maechler M, Bolker B, Walker S. Fitting Linear Mixed-Effects Models Using lme4. Journal of Statistical Software. 2015;67(1):1–48.

24. Oksanen J, Blanchet F, Kindt R. Package, “vegan”. 2015.

25. McMurdie PJ, Holmes S. phyloseq: an R package for reproducible interactive analysis and graphics of microbiome census data. PloS one. 2013;8(4):e61217. PubMed PMID: 23630581. Pubmed Central PMCID: PMC3632530.

26. Segata N, Izard J, Waldron L, Gevers D, Miropolsky L, Garrett WS, et al. Metagenomic biomarker discovery and explanation. Genome Biology. 2011 2011/06/24;12(6):R60.

27. Dice LR. Measures of the Amount of Ecologic Association Between Species. Ecology. 1945;26(3):297–302.

28. Wickham H. ggplot2: Elegant Graphics for Data Analysis. New York, NY: Springer-Verlag; 2016.

29. Schoster A, Kunz T, Lauper M, Graubner C, Schmitt S, Weese JS. Prevalence of Clostridium difficile and Clostridium perfringens in Swiss horses with and without gastrointestinal disease and microbiota composition in relation to Clostridium difficile shedding. Veterinary microbiology. 2019 Dec;239:108433. PubMed PMID: 31767096.

30. Grześkowiak Ł, Dadi TH, Zentek J, Vahjen W. Developing Gut Microbiota Exerts Colonisation Resistance to Clostridium (syn. Clostridioides) difficile in Piglets. Microorganisms. 2019 Jul 26;7(8). PubMed PMID: 31357520. Pubmed Central PMCID: PMC6723027. Epub 2019/07/31. eng.

31. O’Hara E, Kenny DA, McGovern E, Byrne CJ, McCabe MS, Guan LL, et al. Investigating temporal microbial dynamics in the rumen of beef calves raised on two farms during early life. FEMS Microbiology Ecology. 2020;96(2).

32. Haley BJ, Kim S-W, Salaheen S, Hovingh E, Van Kessel JAS. Differences in the Microbial Community and Resistome Structures of Feces from Preweaned Calves and Lactating Dairy Cows in Commercial Dairy Herds. Foodborne Pathogens and Disease. 2020 2020/08/01;17(8):494–503.

33. Gomez DE, Arroyo LG, Costa MC, Viel L, Weese JS. Characterization of the Fecal Bacterial Microbiota of Healthy and Diarrheic Dairy Calves. J Vet Intern Med. 2017;31(3):928–39. PubMed PMID: 28390070. Epub 04/07. eng.

34. Dill-McFarland KA, Breaker JD, Suen G. Microbial succession in the gastrointestinal tract of dairy cows from 2 weeks to first lactation. Sci Rep. 2017;7:40864-. PubMed PMID: 28098248. eng.

35. Henke MT, Kenny DJ, Cassilly CD, Vlamakis H, Xavier RJ, Clardy J. Ruminococcus gnavus, a member of the human gut microbiome associated with Crohn’s disease, produces an inflammatory polysaccharide. Proceedings of the National Academy of Sciences. 2019;116(26):12672.

36. Joossens M, Huys G, Cnockaert M, De Preter V, Verbeke K, Rutgeerts P, et al. Dysbiosis of the faecal microbiota in patients with Crohn’s disease and their unaffected relatives. Gut. 2011 May;60(5):631–7. PubMed PMID: 21209126. Epub 2011/01/07. eng.

37. Hall AB, Yassour M, Sauk J, Garner A, Jiang X, Arthur T, et al. A novel Ruminococcus gnavus clade enriched in inflammatory bowel disease patients. Genome Medicine. 2017 2017/11/28;9(1):103.

38. Chua H-H, Chou H-C, Tung Y-L, Chiang B-L, Liao C-C, Liu H-H, et al. Intestinal Dysbiosis Featuring Abundance of Ruminococcus gnavus Associates With Allergic Diseases in Infants. Gastroenterology. 2018 2018/01/01/;154(1):154–67.

39. Zheng H, Liang H, Wang Y, Miao M, Shi T, Yang F, et al. Altered Gut Microbiota Composition Associated with Eczema in Infants. PloS one. 2016;11(11):e0166026. PubMed PMID: 27812181. Pubmed Central PMCID: PMC5094743. Epub 2016/11/05. eng.

40. Breban M, Tap J, Leboime A, Said-Nahal R, Langella P, Chiocchia G, et al. Faecal microbiota study reveals specific dysbiosis in spondyloarthritis. Annals of the rheumatic diseases. 2017 Sep;76(9):1614–22. PubMed PMID: 28606969. Epub 2017/06/14. eng.

41. Ma T, Villot C, Renaud D, Skidmore A, Chevaux E, Steele M, et al. Linking perturbations to temporal changes in diversity, stability, and compositions of neonatal calf gut microbiota: prediction of diarrhea. The ISME Journal. 2020 2020/09/01;14(9):2223–35.

42. Ling Z, Li Z, Liu X, Cheng Y, Luo Y, Tong X, et al. Altered Fecal Microbiota Composition Associated with Food Allergy in Infants. Applied and Environmental Microbiology. 2014;80(8):2546–54.

43. Zhu JJ, Gao MX, Song XJ, Zhao L, Li YW, Hao ZH. Changes in bacterial diversity and composition in the faeces and colon of weaned piglets after feeding fermented soybean meal. Journal of medical microbiology. 2018 Aug;67(8):1181–90. PubMed PMID: 29923819. Epub 2018/06/21. eng.

44. Youssef O, Lahti L, Kokkola A, Karla T, Tikkanen M, Ehsan H, et al. Stool Microbiota Composition Differs in Patients with Stomach, Colon, and Rectal Neoplasms. Digestive Diseases and Sciences. 2018 2018/11/01;63(11):2950–8.

45. Liang JQ, Li T, Nakatsu G, Chen Y-X, Yau TO, Chu E, et al. A novel faecal Lachnoclostridium marker for the non-invasive diagnosis of colorectal adenoma and cancer. Gut. 2020;69(7):1248.

46. Klein-Jöbstl D, Schornsteiner E, Mann E, Wagner M, Drillich M, Schmitz-Esser S. Pyrosequencing reveals diverse fecal microbiota in Simmental calves during early development. Front Microbiol. 2014 2014-November-17;5(622). English.

47. Malmuthuge N, Griebel PJ, Guan le L. Taxonomic identification of commensal bacteria associated with the mucosa and digesta throughout the gastrointestinal tracts of preweaned calves. Appl Environ Microbiol. 2014 Mar;80(6):2021–8. PubMed PMID: 24441166. Pubmed Central PMCID: 3957634.

48. Tomassini L. Rectal microbiota dynamics in pre-weaned dairy calves depending on colostrum intake, presence of diarrhea and antibiotic treatment: Washington State University; 2015.

49. Hennessy ML, Indugu N, Vecchiarelli B, Bender J, Pappalardo C, Leibstein M, et al. Temporal changes in the fecal bacterial community in Holstein dairy calves from birth through the transition to a solid diet. PloS one. 2020;15(9):e0238882.

50. Han SH, Yi J, Kim JH, Lee S, Moon HW. Composition of gut microbiota in patients with toxigenic Clostridioides (Clostridium) difficile: Comparison between subgroups according to clinical criteria and toxin gene load. PloS one. 2019;14(2):e0212626. PubMed PMID: 30785932.

51. Pérez-Cobas AE, Artacho A, Ott SJ, Moya A, Gosalbes MJ, Latorre A. Structural and functional changes in the gut microbiota associated to Clostridium difficile infection. Front Microbiol. 2014;5:335-. PubMed PMID: 25309515. eng.

52. Eeckhaut V, Machiels K, Perrier C, Romero C, Maes S, Flahou B, et al. Butyricicoccus pullicaecorum in inflammatory bowel disease. Gut. 2013;62(12):1745–52.

53. Devriese S, Eeckhaut V, Geirnaert A, Van den Bossche L, Hindryckx P, Van de Wiele T, et al. Reduced Mucosa-associated Butyricicoccus Activity in Patients with Ulcerative Colitis Correlates with Aberrant Claudin-1 Expression. Journal of Crohn’s and Colitis. 2016;11(2):229–36.

54. Hang BPT, Wredle E, Dicksved J. Analysis of the developing gut microbiota in young dairy calves—impact of colostrum microbiota and gut disturbances. Tropical Animal Health and Production. 2020 2020/12/28;53(1):50.

55. Furusawa Y, Obata Y, Fukuda S, Endo TA, Nakato G, Takahashi D, et al. Commensal microbe-derived butyrate induces the differentiation of colonic regulatory T cells. Nature. 2013 2013/12/01;504(7480):446–50.

56. Alipour MJ, Jalanka J, Pessa-Morikawa T, Kokkonen T, Satokari R, Hynönen U, et al. The composition of the perinatal intestinal microbiota in cattle. Sci Rep. 2018;8(1):10437-. PubMed PMID: 29993024. eng.

